# Metabolomics reveals lipid and amino acid signatures of disease severity in multiple sclerosis

**DOI:** 10.64898/2026.08.04.742797

**Authors:** Rachel E. Rodin, Brain C. Healy, Mariann Polgar-Turcsanyi, Hrishikesh A. Lokhande, Tanuja Chitnis

## Abstract

**Objective:** Plasma metabolomics offers insight into multiple sclerosis (MS) pathophysiology, but existing studies are limited by small sample sizes and incomplete clinical data.

**Methods:** We conducted plasma metabolomic profiling of 411 deeply phenotyped patients with MS and 46,443 controls, analyzing 162 metabolites in 30 biologically related metabolite groups. We characterized associations with MS diagnosis, disability, disease subtype, and inflammatory disease activity using regression and differential network enrichment analysis. We additionally examined 25 pre-diagnosis individuals whose samples were collected before their first demyelinating event.

**Results:** Fourteen of 30 metabolite groups were associated with MS after false discovery rate correction, with the strongest positive associations observed for atherogenic lipoproteins, glycine, cholines, and saturated fatty acids, and the strongest negative associations for aromatic amino acids, branched-chain amino acids, alanine, and citrate. Differential network enrichment analysis identified two dysregulated subnetworks encompassing amino acid and energy metabolism and lipid and lipoprotein metabolism. Five metabolite groups were negatively associated with disability: small high-density lipoprotein particles, histidine, branched-chain amino acids, albumin, and aromatic amino acids. The omega-6/omega-3 fatty acid ratio was significantly associated with recent relapse (*OR* = 1.92, FDR-*p* = 0.030) and nominally associated with future MRI activity, especially in patients on moderate or high-efficacy disease-modifying therapy. The MS metabolic signature was not detectable in pre-diagnosis samples.

**Interpretation:** These findings highlight coordinated dysregulation of amino acid and lipoprotein metabolism as hallmarks of established MS and identify a novel association of the omega-6/omega-3 ratio with inflammatory disease activity.

## Introduction

Multiple sclerosis (MS) is a chronic immune-mediated demyelinating disease of the central nervous system affecting approximately 2.8 million people worldwide.^1^ Despite advances in disease-modifying therapies that dramatically reduce relapse rates, many patients continue to accumulate disability.^2, 3^ Reliable biomarkers associated with diagnosis, disease activity, and disability remain an important unmet need for clinical care and mechanistic research.^4^

Metabolomics offers a window into the biological processes underlying MS.^5^ Prior studies have identified abnormalities in blood, cerebrospinal fluid, and stool metabolites, including alterations in amino acids, lipids, and lipoproteins.^6^ Several reports have described reduced aromatic amino acids, branched-chain amino acids, and arginine in blood of patients with MS, while others have identified abnormalities in fatty acids, sphingolipids, and lipoprotein subclasses.^7–14^ Some studies suggest that altered amino acid and lipid profiles correlate with disease severity and can potentially aid in predicting the disease course.^7, 12, 15, 16^ However, many studies have been limited by small sample sizes, heterogeneous metabolomics platforms, and incomplete clinical phenotyping, and several key findings have not been consistently replicated including contradictory findings regarding amino acids in MS.^14, 15^

Many questions remain unanswered in MS metabolomics. First, the extent to which plasma metabolomic dysregulation in MS reflects disease-specific pathophysiology versus comorbid conditions or treatment effects has not been rigorously evaluated in a large, covariate-adjusted cohort. Second, the respective metabolomic correlates of demyelinating relapses and disability progression have not been comprehensively assessed. Third, whether metabolomic perturbations are detectable prior to the first clinical demyelinating event remains unknown.

To address these questions, we performed plasma metabolomic profiling in a large set of deeply phenotyped patients with MS and controls. We implemented comprehensive regression analyses, differential network enrichment analysis, and longitudinal disease activity analyses to characterize the metabolome across the spectrum of MS clinical phenotypes. We additionally examined a unique cohort of 25 individuals whose samples were collected prior to their first demyelinating event to evaluate whether metabolomic changes precede clinical disease onset.

## Materials and Methods

### Study population and metabolomic profiling

#### Participants and biosamples

The Mass General Brigham biobank has enrolled more than 135,000 participants across clinical sites in greater Boston, United States since 2008.^17^ Participant biosamples are linked to electronic health record data and self-reported health surveys.^18^ Blood samples were collected by venipuncture, processed, and stored at −80°C until analysis. All participants provided written informed consent. The Human Research Committee of Mass General Brigham approved the Biobank research protocol (IRB protocol 2009P002312). This study was conducted in accordance with the Declaration of Helsinki.

#### Metabolomics data

Banked plasma samples from 47,194 biobank participants underwent metabolomic profiling using the Nightingale Health nuclear magnetic resonance platform, which has previously been used in other large population-based biobanks.^19^ Quality control was performed similarly to previously published methods (Supplementary Methods).^20^ After filtering to minimize redundancy, 162 unique metabolites (Supplementary Table 1) were used for downstream analyses.

#### Identification of control group

Samples with >20% missingness and participants under age 18 were excluded. Participants with a self-reported diagnosis of MS not confirmed on electronic health record review were excluded. Lastly, 25 patients who developed MS after metabolomics sampling were excluded from the control group. In total, 46,443 participants without MS were classified as controls. Race, ethnicity, sex, age, and Charlson comorbidity index data were obtained from the biobank.^21^

#### Clinical phenotyping of multiple sclerosis cases

Of 932 participants initially identified as having MS, 411 patients met 2024 McDonald Criteria for MS and had a clinical visit within one year before or after sample collection. We included participants who had a diagnosis of clinically isolated syndrome (CIS) at the time of sampling and later developed MS. Remaining participants were either classified as pre-diagnosis MS (n = 25) or excluded due to lack of neurology clinical data or not meeting diagnostic criteria. Biosampling occurred between 2011 and 2023 for MS cases, with a median sampling year of 2017. Approximately half of the 411 participants with MS were seen at the Brigham and Women’s Hospital Multiple Sclerosis Center with clinical data documented by neurologists in an Oracle-based database. Clinical data for remaining participants was extracted from the medical record by an MS specialist. Patients were classified as having CIS, relapsing remitting multiple sclerosis (RRMS), secondary progressive multiple sclerosis (SPMS), or primary progressive multiple sclerosis (PPMS) at the visit closest to biobank sample collection. The Expanded Disability Status Scale (EDSS) score was extracted directly from a clinical neurology note or derived from a neurologic exam and history using an EDSS calculation tool.^22^ Relapse, MRI, and disease-modifying therapy (DMT) efficacy were ascertained as detailed in Supplementary Methods.

### Statistical analysis

Analyses were performed in R version 4.5.3 using packages including DNEA (version 3.22), ggplot2, and emmeans. Metabolite values were imputed, grouped into 30 biologically informed clusters by hierarchical clustering of Spearman correlations, and analyzed as mean z-scores per group (Supplementary Methods, Supplementary Tables 1 and 2, Supplementary Figure 1). In analyses of clinical outcomes, metabolites or metabolite group z-scores were the predictor; in diagnostic group comparisons, disease status was the predictor. False discovery rate (FDR) correction was applied to all analyses using the Benjamini-Hochberg method.

#### Identifying a metabolic signature of multiple sclerosis

Linear regression was used to estimate the difference in mean metabolite z-scores comparing 411 MS cases to 46,443 controls, adjusting for age, sex, race (Caucasian or non-Caucasian), ethnicity (Hispanic or non-Hispanic), and Charlson comorbidity index. Regressions were completed using all 162 metabolites separately (Fig. 1A, Supplementary Table 3) and in 30 statistically clustered groups (Fig. 1B, Supplementary Table 4).

**Figure 1.**
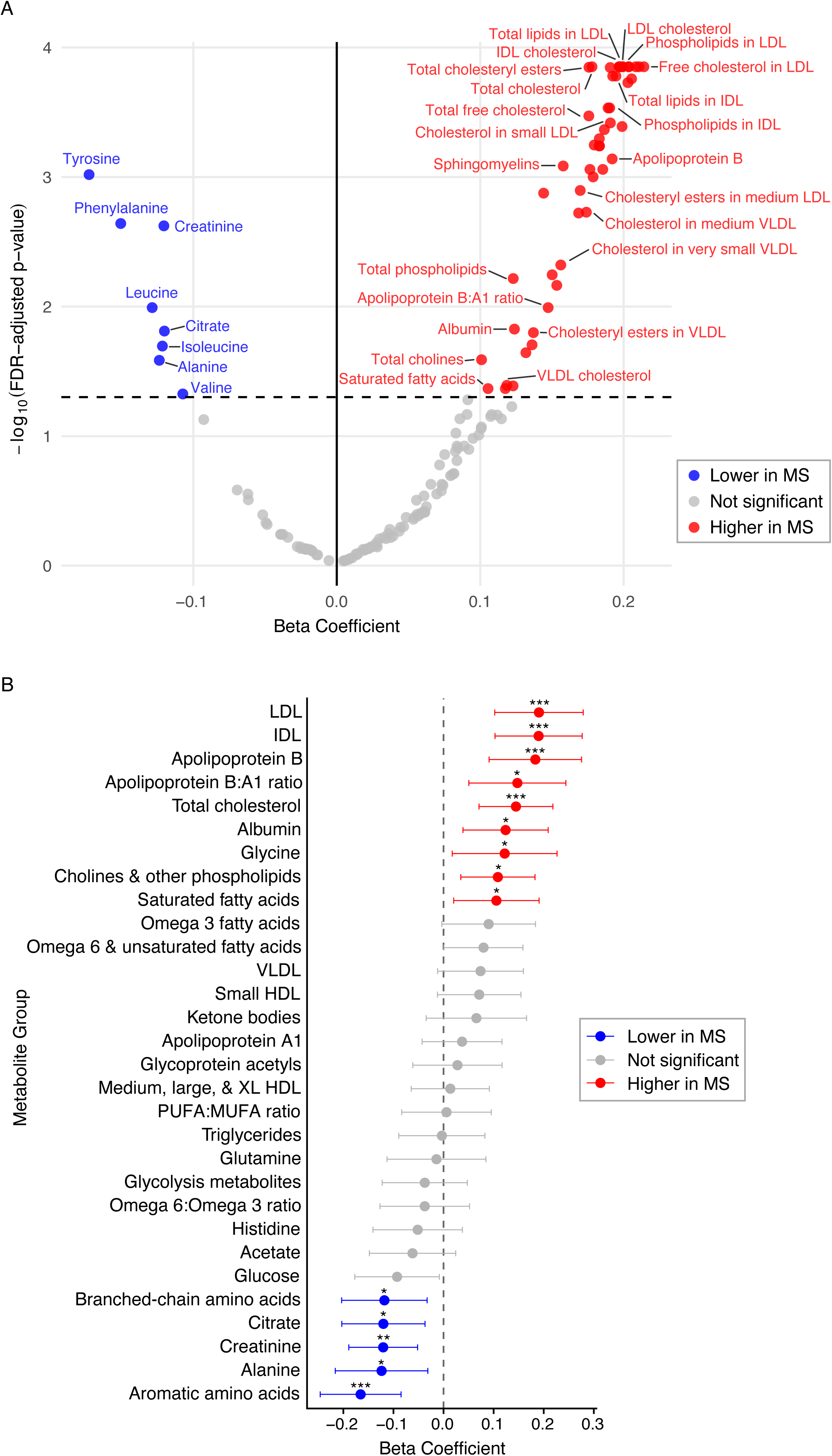
Metabolomic profiles of patients with multiple sclerosis versus controls. (A) Volcano plot showing differences in 162 metabolites between people with MS and controls, with the x-axis representing linear regression coefficients (change in metabolite z-score) and the y-axis representing −log10(*FDR-p-*value) from the regression. Colored dots represent FDR < 0.05. Some key metabolites are labeled. (B) Forest plot showing differences in 30 metabolite groups between people with MS and controls, with the x-axis again representing linear regression coefficients (change in metabolite z-score). Colored dots represent FDR-p < 0.05. Error bars represent 95% confidence interval. Asterisks indicate statistical significance. *FDR-p < 0.05, **FDR-p < 0.01, ***FDR-p < 0.001.

#### Differential network enrichment analysis

Differential Network Enrichment Analysis (DNEA) was performed to identify dysregulated metabolic subnetworks between cases and controls using the DNEA R package, with subnetwork significance assessed by the Network-level Gene Set Analysis (NetGSA) and FDR correction across subnetworks (Supplementary Methods, Supplementary Tables 5-7). A 1:10 subsampling sensitivity analysis confirmed robustness of findings to group size imbalance (Supplementary Table 8).

#### Analysis of disability

Linear regression was used to assess associations between the 30 metabolite groups and EDSS score at the visit closest to sampling, adjusting for age, sex, race, BMI, and DMT efficacy (Supplementary Table 9).

#### Disease subtype and progression

Logistic regression was used to compare metabolite levels between progressive (PPMS + SPMS; *n* = 85) and relapsing (RRMS + CIS; *n* = 326) MS patients, adjusting for age, sex, race, BMI, and DMT efficacy (Supplementary Table 10). Linear regression was used to compare metabolite levels across MS subtypes (CIS, PPMS, SPMS) with RRMS as the reference group, adjusting for age, sex, race, BMI, EDSS, and DMT efficacy (Supplementary Fig. 2, Supplementary Table 11). Cox regression was used to assess time to secondary progressive MS diagnosis among RRMS patients who later converted to SPMS (*n* = 38), adjusting for age, sex, BMI, DMT efficacy, and baseline EDSS (Supplementary Table 12).

#### Inflammatory disease activity

Logistic regression was used to assess associations between metabolite groups and four binary disease activity outcomes: relapse in the 12 months before sample, relapse in the 12 months after sample, MRI activity in the 12 months before sample, and MRI activity in the 12 months after sample. Covariates were age, sex, race, EDSS, DMT efficacy, and BMI. FDR correction was applied across 30 metabolite groups within each outcome (Fig. 4A-B, Supplementary Table 13). In post-hoc analyses, the omega-6/omega-3 fatty acid ratio was examined across six disease activity outcomes with FDR correction across outcomes (Supplementary Table 14), and its interaction with DMT efficacy (none, low, moderate/high) was assessed using a multiplicative interaction term with stratified analyses (Supplementary Table 15).

#### MRI and blood biomarkers

A subset of 28 patients had serum neurofilament light chain (NfL) and 26 had glial fibrillary acidic protein (GFAP) measurements from within one year of metabolomics sampling. Additionally, a subset of 130 patients had 3-Tesla brain MRIs within one year of the metabolomics sample, with T2-lesion volume and brain parenchymal fraction calculated according to previously published methods.^23, 24^ Linear regression was used to assess associations between metabolite groups and NfL, GFAP, T2-lesion volume, and brain parenchymal fraction, adjusting for race, age, sex, BMI, and DMT efficacy (Supplementary Table 16).^23, 24^

#### Pre-diagnosis multiple sclerosis

Twenty-five participants had not yet experienced a demyelinating attack at the time of sample collection but later went on to be diagnosed with MS (Supplementary Table 17). Linear regression was used to compare pre-diagnosis patients and established MS patients to controls, adjusting for age, sex, and race, with controls as the reference group. Pre-diagnosis patients were next compared to established MS patients using the same parameters, with established MS patients as the reference group (Supplementary Table 18). To visualize covariate-adjusted metabolite levels across all three groups, analysis of covariance (ANCOVA) was performed across all MS, control and pre-diagnosis participants, adjusting for age, sex, and race, with controls as the reference group (Supplementary Figs. 3 and 4).

## Results

### Study population

Among Mass General Brigham Biobank participants who underwent plasma metabolomic profiling, 411 had MS or CIS at the time of sample collection and 46,443 were controls. The MS cohort was predominately RRMS, had a mean age of 49.4 years, and was majority female and White. Approximately 71% of patients were on a disease-modifying therapy at the time of sampling. Controls were older on average (mean age 54.0 years) with a higher proportion of males (Table 1).

**Table 1.** Summary characteristics of study population at time of plasma collection.

| Diagnosis | MS | Control |
| --- | --- | --- |
| Number of participants | 411 | 46,443 |
| Age, years, mean (SD) | 49.4 (13.0) | 54.0 (16.6) |
| Age range (min, max) | 18.9, 81.8 | 18.0, 95.8 |
| Sex, female, *n* (%) | 301 (73.2%) | 25,274 (54.4%) |
| Race, *n* (%) |  |  |
| White | 365 (88.8%) | 39,411 (84.9%) |
| Black | 20 (4.9%) | 2186 (4.7%) |
| Asian/Pacific Islander | 2 (5.8%) | 1154 (2.5%) |
| Multiple/Other/Unknown | 24 (5.8%) | 3692 (7.9%) |
| Ethnicity, *n* (%) |  |  |
| Non-Hispanic | 364 (88.6%) | 40,844 (87.9%) |
| Hispanic | 8 (1.9%) | 1326 (2.9%) |
| Other/Unknown | 39 (9.5%) | 4273 (9.2%) |
| Charlson Index, mean (SD) | 3.9 (2.6) | 4.6 (2.6) |
| BMI, kg/m<sup>2</sup>, mean (SD) | 28.4 (6.5) |  |
| BMI, kg/m<sup>2</sup>, range (min, max) | 17.0, 53.2 |  |
| EDSS, mean (IQR) | 2.7 (1.5 – 3.5) |  |
| EDSS, range (min, max) | 0, 9.0 |  |
| Follow-Up Interval, years, mean (SD) | 6.3 (3.3) |  |
| MS Diagnosis, *n* (%) |  |  |
| CIS | 9 (2.2%) |  |
| RRMS | 317 (77.1%) |  |
| SPMS | 63 (15.3%) |  |
| PPMS | 22 (5.4%) |  |
| DMT efficacy class, *n* (%) |  |  |
| High Efficacy | 57 (13.9%) |  |
| Moderate Efficacy | 117 (28.4%) |  |
| Low Efficacy | 118 (28.7%) |  |
| None | 119 (29.0%) |  |
| Relapse or New T2/T1Gd+ within ±1 year, *n* (%) | 112 (27.3%) |  |

### Metabolomic signature of multiple sclerosis

We identified 59 individual metabolites significantly associated with MS after FDR correction (Supplementary Table 3, Fig. 1A). At the metabolite group level, 14 of 30 metabolite groups were significantly associated with MS after FDR correction (Fig. 1B, Supplementary Fig. 1, Supplementary Table 4). The strongest positive associations with MS were observed for atherogenic lipoprotein pathways including intermediate-density lipoprotein (IDL; β = 0.190, FDR-*p* = 0.0003), low-density lipoprotein (LDL; β = 0.190, FDR-*p* = 0.0003), total cholesterol (β = 0.144, FDR-*p* = 0.0007), and apolipoprotein B (β = 0.183, FDR-*p* = 0.0007). Conversely, aromatic amino acids (β = −0.166, FDR-*p* = 0.0006), branched-chain amino acids (β = −0.118, FDR-*p* = 0.018), creatinine (β = −0.121, FDR-*p* = 0.003), citrate (β = −0.120, FDR-*p* = 0.014), and alanine (β = −0.124, FDR-*p* = 0.022) were significantly lower in MS patients compared to controls. Cholines and other phospholipids (β = 0.108, FDR-*p* = 0.014), saturated fatty acids (β = 0.105, FDR-*p* = 0.035), glycine (β = 0.122, FDR-*p* = 0.048), and albumin (β = 0.124, FDR-*p* = 0.014) were also significantly elevated in MS (Fig. 1B, Supplementary Table 4).

### Differential network enrichment analysis

To characterize the systems-level organization of metabolic dysregulation in MS, DNEA was performed on residualized metabolite values across MS cases and controls. Two significantly dysregulated metabolic subnetworks were identified (Fig. 2A, Supplementary Tables 5-7).

**Figure 2.**
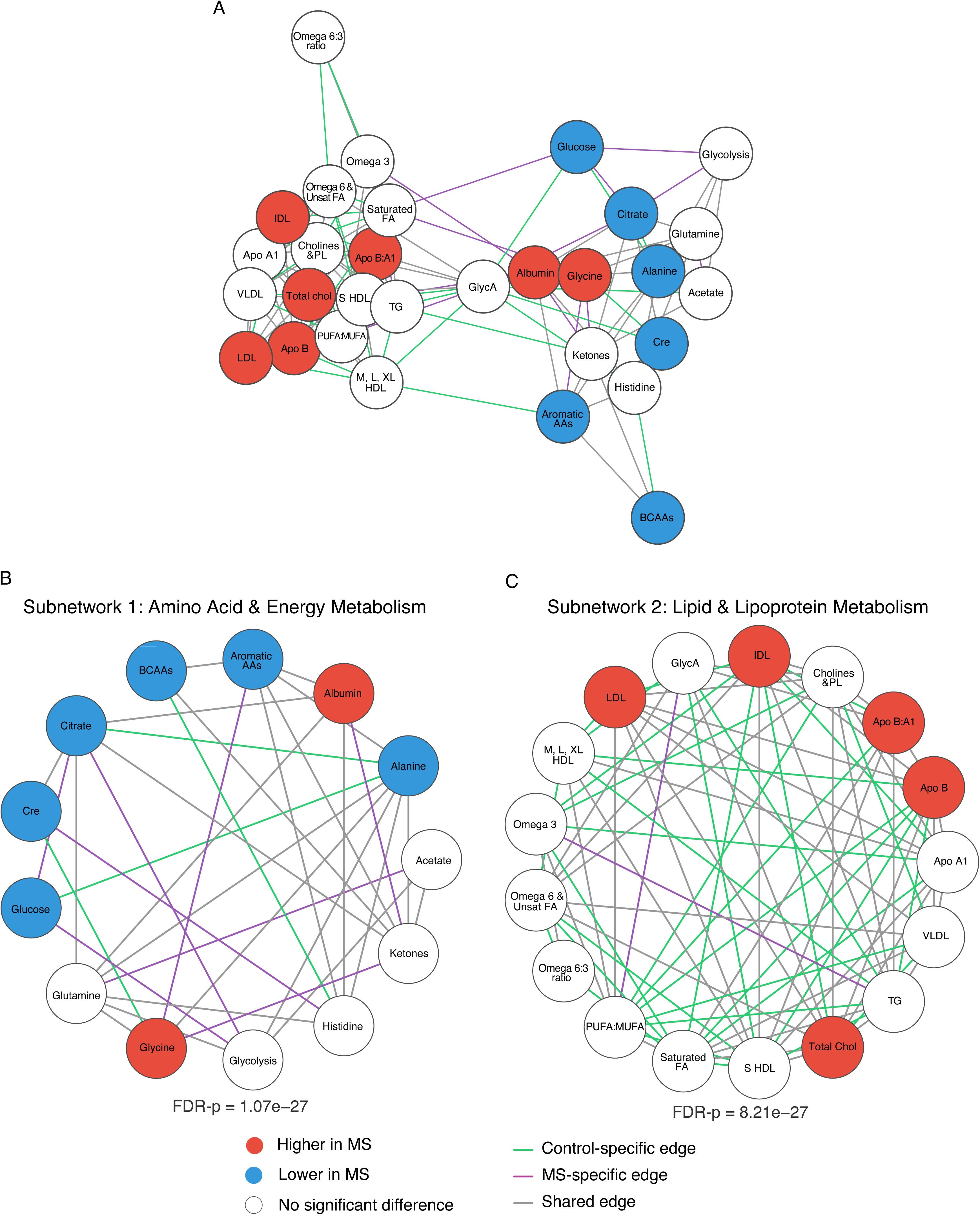
Differential network enrichment analysis of metabolite groups in multiple sclerosis versus controls. (A) Overall network analysis of all 30 metabolite groups in MS versus controls. (B) Subnetwork 1, representing amino acid and energy metabolism. (C) Subnetwork 2, representing lipid and lipoprotein metabolism. In all panels circles represent nodes (metabolite groups) and lines represent edges (associations between metabolites). Statistical significance is based on a network-level Gene Set Analysis test, with false discovery rate correction applied across subnetworks.

Subnetwork 1 encompassed amino acid and energy metabolism (13 nodes, 34 edges), with 8 of 13 nodes differentially expressed after FDR correction (NetGSA *p* = 5.33×10 ², FDR-*p* = 1.07×10 ²). Differentially expressed nodes were predominantly depleted in MS (aromatic amino acids, creatinine, citrate, branched-chain amino acids, alanine, and glucose), while albumin and glycine were significantly elevated in MS (Supplementary Table 5). Beyond differential expression of individual nodes, the network topology was reorganized in MS (Fig. 2B). Several metabolic connections present in controls were lost in MS, including alanine-glucose, alanine-citrate, and creatinine-glycine co-regulation, suggesting disruption of normal amino acid-energy coupling. Conversely, new MS-specific edges emerged, including citrate-glucose, glucose-glycolysis metabolites, and novel connections from albumin to ketone bodies and small high-density lipoprotein (HDL), indicating disease-associated alterations in energy metabolism (Supplementary Table 6).

Subnetwork 2 encompassed lipid and lipoprotein metabolism (17 nodes, 81 edges), with 5 of 17 nodes differentially expressed, all of which were upregulated in MS (LDL, IDL, apolipoprotein B, apolipoprotein B to A1 ratio, and total cholesterol; NetGSA *p* = FDR-*p* = 8.21×10 ²) (Fig. 2C). The lipid subnetwork showed extensive reorganization in MS, with multiple connections present in controls and lost in MS, including apolipoprotein B connections to saturated fatty acids, small HDL, triglycerides, and medium/large HDL, as well as the IDL-LDL coregulation edge (Supplementary Table 6). This decoupling of normally coordinated lipoprotein relationships suggests that the elevation of atherogenic lipoproteins in MS occurs in the context of broader disruption of lipoprotein network architecture rather than a uniform upregulation of lipid molecules. Sensitivity analysis using a 1:10 subsampled dataset confirmed robustness of both subnetworks to group size imbalance (Supplementary Table 8).

### Metabolic associations with disability

We identified five metabolite groups significantly associated with EDSS at the time of blood sampling after FDR correction (Fig. 3, Supplementary Table 9). Small HDL (β = −0.381, FDR-*p* = 0.006), histidine (β = −0.328, FDR-*p* = 0.008), branched-chain amino acids (β = −0.338, FDR-*p* = 0.008), albumin (β = −0.343, FDR-*p* = 0.008), and aromatic amino acids (β = −0.329, FDR-*p* = 0.016) were negatively associated with EDSS. No metabolite group was positively associated with EDSS after FDR correction, though the general inflammatory marker glycoprotein acetyls, previously associated with progressive MS in one study, was nominally positively correlated with EDSS (Supplementary Table 9).

**Figure 3.**
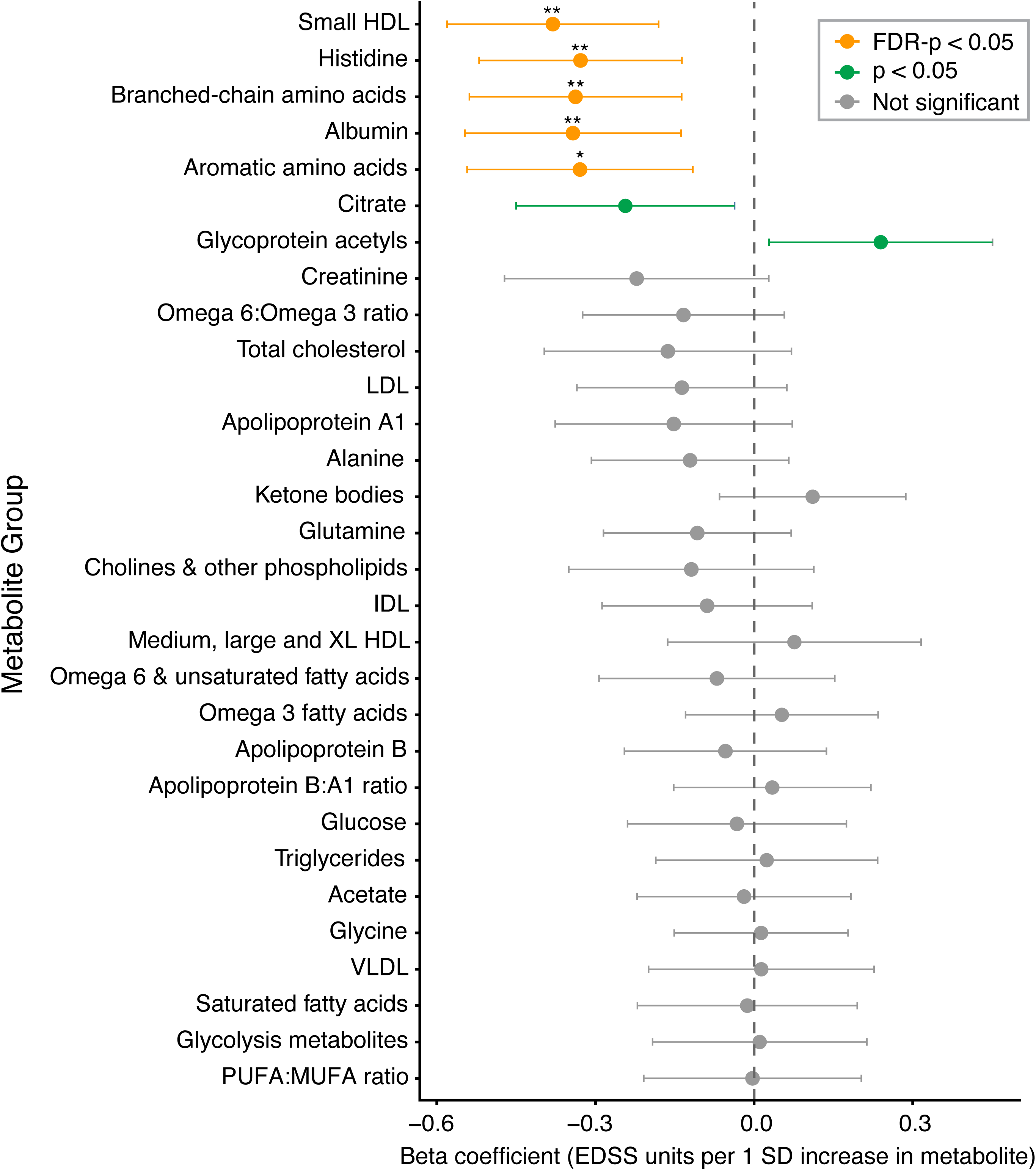
Association of metabolite groups with Expanded Disability Status Scale (EDSS) score in patients with multiple sclerosis. Forest plot showing relationship between the 30 metabolite groups and EDSS. The x-axis represents coefficients from the linear regression, or EDSS units per one standard deviation change in metabolite value. Negative values indicate an inverse relationship between metabolite level and EDSS. Error bars represent 95% confidence interval. Asterisks above orange dots indicate statistical significance. *FDR-p < 0.05, **FDR-p < 0.01, ***FDR-p < 0.001. Green dots represent nominal significance (*p* < 0.05) that did not survive FDR correction.

### Metabolic differences by disease subtype and disability progression

Logistic regression comparing progressive (PPMS or SPMS) to relapsing (RRMS or CIS) patients identified no metabolite groups significantly associated with MS progression after FDR correction (Supplementary Table 10). Linear regression comparing all MS subtypes to RRMS as the reference group likewise identified no FDR-significant associations for SPMS, PPMS, or CIS, though CIS had nominally higher lipids (LDL, IDL, and apolipoprotein B) and PPMS had nominally higher aromatic amino acids and lower omega-6/omega-3 ratio compared to RRMS (Supplementary Table 11, Supplementary Fig. 2). Cox regression assessing time to SPMS conversion among 300 RRMS patients with available follow-up data, of whom 38 converted to SPMS over a mean follow-up of 6.2 years, identified no metabolite group significantly associated with conversion after FDR correction (Supplementary Table 12). Both aromatic amino acids and branched-chain amino acids showed nominally lower hazard of SPMS conversion, consistent with the direction of their EDSS associations, but neither association survived FDR correction (Supplementary Table 12). Together, these findings suggest that the metabolomic differences observed in MS primarily reflect current disability severity rather than disease subtype or future disease trajectory.

### Omega 6/Omega 3 ratio and inflammatory disease activity

Among 399 MS cases with available relapse and MRI data, logistic regression was used to identify metabolite groups significantly associated with relapse in the prior 12 months, MRI activity in the prior 12 months, relapse in the following 12 months, or MRI activity in the following 12 months (Supplementary Table 13). The omega-6/omega-3 ratio was significantly associated with relapse in the prior 12 months (*OR* = 1.92, FDR-*p* = 0.030; Fig. 4A) and nominally associated with MRI activity in the following 12 months (*OR* = 1.98, FDR-*p* = 0.059; Fig. 4B). Given this finding, post-hoc hypothesis-generating analyses were performed to more comprehensively characterize the relationship between the omega-6/omega-3 ratio and disease activity. These analyses were not pre-specified and should be interpreted as exploratory. The omega-6/omega-3 ratio was examined across six disease activity outcomes, and a higher omega-6/omega-3 ratio was significantly associated with prior relapse (*OR* = 1.94), future MRI activity (*OR* = 1.97) and prior relapse or MRI activity (*OR* = 1.66, all FDR-*p* = 0.005) when FDR-corrected across six outcomes (Fig. 4C, Supplementary Table 14).

**Figure 4.**
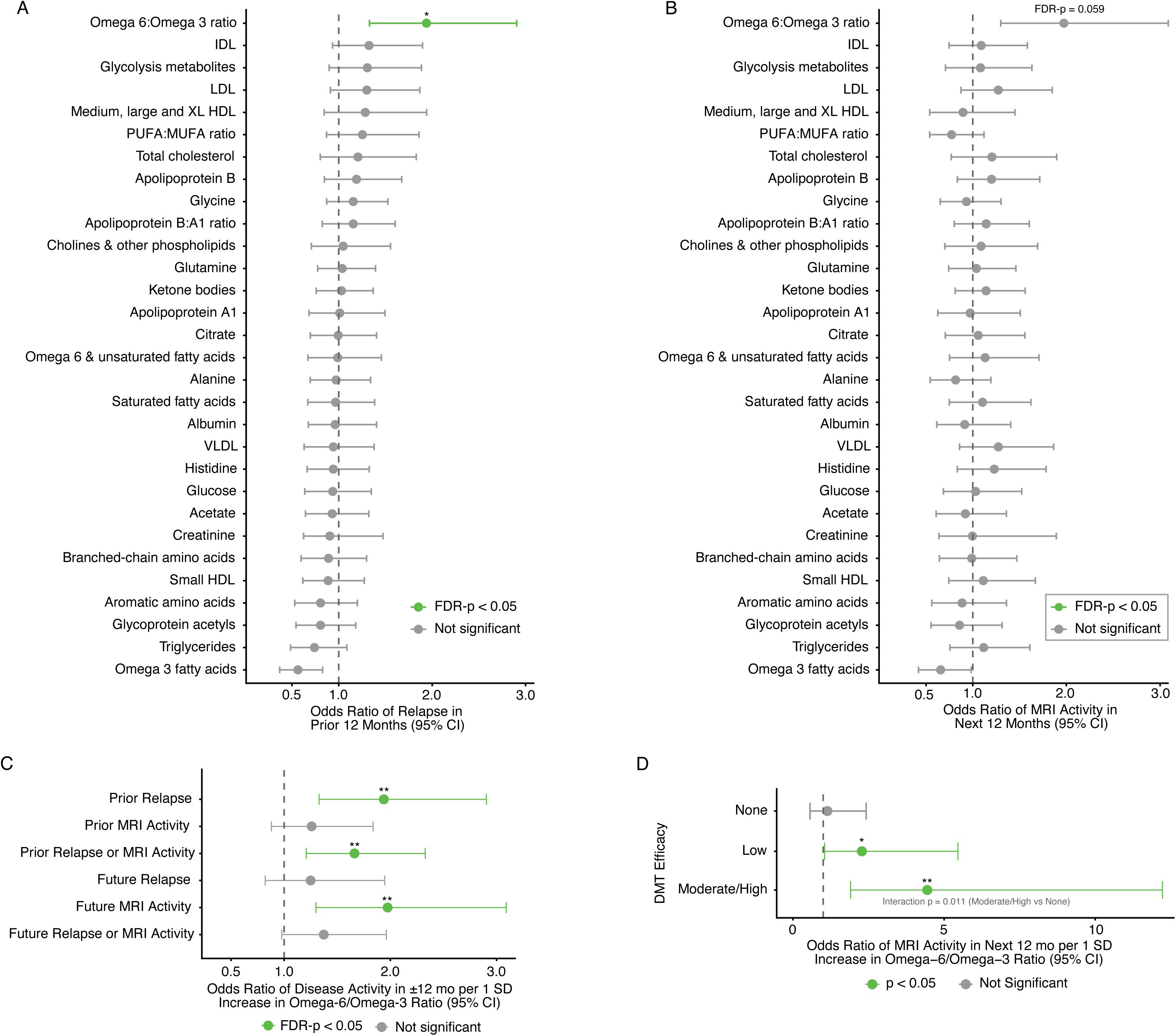
The ratio of omega-6 to omega-3 fatty acids corresponds to past and future multiple sclerosis disease activity. (A) The forest plot shows results of a logistic regression for all 30 metabolite groups and relapse in the 12 months preceding sample collection. (B) The forest plot shows results of a logistic regression for all 30 metabolite groups and MRI activity in the 12 months following sample collection. (C) Logistic regression for the omega-6/omega-3 ratio and six disease activity outcomes. (D) Logistic regression for omega-6/omega-3 ratio and MRI activity in the 12 months following sample collection, separated by DMT efficacy (none, low, or moderate/high). The interaction between the omega-6/omega-3 ratio and DMT efficacy was significant at *p* = 0.011 for moderate/high-efficacy versus no DMT. For all panels, the x-axis represents odds ratio and error bars represent 95% confidence interval. Asterisks indicate statistical significance. *FDR-p < 0.05, **FDR-p < 0.01, ***FDR-p < 0.001.

The interaction between the omega-6/omega-3 ratio and DMT efficacy was examined in a post-hoc exploratory analysis prompted by the primary disease activity findings. A significant interaction was identified between the omega-6/omega-3 ratio and DMT efficacy for future MRI activity (likelihood ratio test *p* = 0.035, Fig. 4D, Supplementary Table 15). Stratified analyses demonstrated a marked dose-dependent amplification of the association with increasing DMT efficacy: the omega-6/omega-3 ratio was not significantly associated with future MRI activity among patients on no DMT (*OR* = 1.21, *p* = 0.59), showed a significant association among patients on low-efficacy DMT (*OR* = 2.28, *p* = 0.047), and a markedly stronger association among patients on moderate- or high-efficacy DMT (*OR* = 4.34, *p* = 0.002). No significant interaction with DMT efficacy was observed for other disease activity outcomes.

### MRI and blood biomarker investigations

In a subset of 130 MS patients with quantitative 3T brain MRI data from within one year of metabolomic sampling, no metabolite group was significantly associated with T2 lesion volume or brain parenchymal fraction after FDR correction (Supplementary Table 16). Aromatic amino acids showed a nominally significant inverse association with log-transformed T2 lesion volume, consistent with the direction of their EDSS association. In subsets of 28 and 26 patients with available serum NfL and GFAP measurements respectively, no metabolite group was significantly associated with either biomarker after FDR correction, likely reflecting limited statistical power.

### Pre-diagnosis multiple sclerosis metabolomic profiling

To determine whether the metabolomic signature of MS precedes the first clinical demyelinating event, linear regression was used to compare metabolite levels in 25 pre-diagnosis participants to all controls and established MS patients. Characteristics of the pre-diagnosis cohort are detailed in Supplementary Table 17. The median time to MS diagnosis was 3.7 years (range 0.07–11.4 years). No metabolite group was significantly associated with pre-diagnosis MS status compared to controls or established MS patients after FDR correction (Supplementary Table 18, Supplementary Figs. 3 and 4). Numerically, pre-diagnosis participants showed intermediate metabolite levels between controls and MS patients for several metabolite groups with established MS associations.

## Discussion

In this large cohort, we identified a comprehensive plasma metabolomic signature of MS characterized by two major axes of dysregulation: reduced metabolites of amino acid and energy metabolism, and elevated atherogenic lipoprotein fractions. Within the MS cohort, altered amino acid and lipid profiles were significantly associated with disability severity and inflammatory disease activity. In an analysis of pre-diagnosis cases, these metabolomic differences were not detectable in the years preceding clinical disease onset, suggesting that they reflect established disease processes rather than a MS susceptibility state.

### Amino acid dysregulation in multiple sclerosis

Across our analyses we consistently identified downregulation of aromatic and branched-chain amino acids as a hallmark of MS. Both groups of amino acids were significantly reduced compared to controls, negatively associated with EDSS, and formed the most significantly enriched subnetwork in the DNEA. Additionally, there were trends toward lower branched-chain and aromatic amino acid levels predicting time to SPMS conversion, and lower aromatic amino acids nominally correlated with higher T2 lesion volume on brain MRI. These findings expand on several prior studies and firmly establish aromatic and branched-chain amino acid depletion as core markers of the MS disease state. We also show that the non-essential amino acid alanine is significantly reduced in MS, in contrast to a small study that identified higher alanine in elderly patients with SPMS.^8^ While histidine was not significantly decreased in our overall cohort of MS compared to controls, it did significantly inversely correlate with EDSS. This novel finding is complementary to a report of lower histidine correlating with increased fatigue in female patients with MS.^25^

Whether reduction in circulating amino acids contributes directly to MS pathogenesis or reflects a consequence of disease-associated metabolic changes remains unclear. Aromatic amino acids are known to be involved in remyelination of the central nervous system and may play a role in regulating microglial activation.^26, 27^ Branched-chain amino acids impact microglial function and promote T-regulatory cell survival.^28–30^ Amino acid changes in MS could be mediated through diet, mitochondrial dysfunction, or the gut microbiome.^31, 32^ Our small analysis of 25 pre-diagnosis patients suggests that lower amino acids are not genetically pre-determined in patients who later develop MS. There is growing interest in amino acid supplementation in patients with MS, but more research is needed to determine whether exogenously altering amino acid metabolism can impact the disease course.^33, 34^

Our analysis also revealed extensive rewiring of amino acid and energy metabolism networks in MS. Loss of normal alanine-glucose, alanine-citrate, and branched-chain amino acid-histidine co-regulation, alongside emergence of new disease-specific connections including aromatic amino acids and glycine, suggests a generalized shift in amino acid and energy substrate utilization. These network-level changes may be related to mitochondrial dysfunction and altered immune cell metabolism.^35, 36^

### Lipoprotein dysregulation in multiple sclerosis

The elevation of LDL, IDL, total cholesterol, sphingomyelins, cholines, phospholipids, and apolipoprotein B identified in our MS cohort, and their coordinated upregulation in our network analysis, is consistent with other reports implicating lipoprotein dysregulation in MS pathophysiology.^37–39^ Dysregulated lipid metabolism may contribute to MS pathogenesis through altered T-cell function.^40^ Notably, DNEA revealed that the elevation of atherogenic lipoproteins in MS occurs in the context of extensive network decoupling, implying a fundamentally altered lipoprotein metabolic state rather than simple upregulation of cholesterol synthesis pathways.

While numerous atherogenic lipids associated with MS diagnosis in our study, none were associated with EDSS except small HDL, which negatively correlated with EDSS. The inverse correlation between small HDL and EDSS likely reflects the neuroprotective and anti-inflammatory functions of small HDL particles, which are efficient mediators of ABCA1-dependent cholesterol efflux – a process critical for clearing myelin-derived cholesterol debris and resolving neuroinflammation – and have been independently associated with greater grey matter volume in population-based studies.^41, 42^ Consistent with this, higher HDL is associated with reduced blood-brain barrier injury and decreased pro-inflammatory cell extravasation into cerebrospinal fluid after a first demyelinating event, supporting a direct link between functional small HDL particles and lower disability in MS.^43^

### Omega-6/omega-3 ratio and inflammatory disease activity

The novel finding that elevated omega-6/omega-3 fatty acid ratio corresponds to recent relapse and future MRI activity in patients with MS aligns with converging evidence from observational, mechanistic, and interventional studies. Omega-6 polyunsaturated fatty acids, particularly arachidonic acid, serve as substrates for pro-inflammatory eicosanoids that are linked to MS pathogenesis. In a population-based MS cohort, elevated levels of the arachidonic acid derivative 15-HETE correlated with higher EDSS, elevated serum NfL, lower brain volume, and greater lesion volume.^11^ Conversely, omega-3 polyunsaturated fatty acids (including fish-derived docosahexaenoic acid and plant-derived alpha-linolenic acid) generate anti-inflammatory mediators, inhibit Th1/Th17 polarization, and promote anti-inflammatory microglial phenotype switching.^44, 45^ In the BENEFIT trial, higher baseline serum alpha-linolenic acid was associated with significant reduction in relapse risk.^46^ Thus, a higher omega-6/omega-3 ratio likely reflects a systemic pro-inflammatory lipid milieu that both marks recent disease activity and sustains the neuroinflammatory substrate for future MRI lesion development.

The omega-6/omega-3 ratio may capture the net balance between these opposing lipid pathways more informatively than either fatty acid class alone.^11, 47^ A dietary shift toward higher omega-6 consumption has been proposed to promote chronic low-grade systemic inflammation that may lower the threshold for autoimmune disease flares. While randomized trials of omega-3 supplementation in RRMS have not demonstrated a significant difference in relapse rates or MRI activity, these trials were limited by small sample sizes and notably used omega-6 oils as the comparator, potentially obscuring a true effect by providing an active control rather than inert placebo.^48^ The present finding that the ratio itself, rather than absolute levels of either fatty acid class, predicts both retrospective clinical and prospective radiographic disease activity supports the concept that the relative balance between pro-inflammatory and anti-inflammatory lipid mediator pathways is a meaningful biomarker of the MS inflammatory burden.

The observation that a higher omega-6/omega-3 ratio predicts relapse and MRI activity preferentially in patients on high/moderate-efficacy DMTs – but not in untreated patients – likely reflects the fact that DMTs suppress the dominant adaptive immune drivers of MS relapse, thereby unmasking the contribution of residual innate and lipid-mediated inflammatory pathways that are not directly targeted by these therapies.^49^ Arachidonic acid-derived lipid mediators are produced primarily by myeloid cells and microglia within MS lesions, and their biosynthesis is enhanced by a higher omega-6/omega-3 substrate ratio but is largely unaffected by lymphocyte-targeting DMTs.^11, 50^ In untreated patients, the overwhelming variance in disease activity attributable to uncontrolled adaptive immunity obscures the comparatively modest signal from the omega-6/omega-3 balance, whereas in DMT-treated patients, this lipid-mediated inflammatory pathway becomes a detectable determinant of breakthrough disease activity. These findings suggest that optimizing the omega-6/omega-3 ratio may represent a complementary strategy in DMT-treated patients.

### Pre-diagnosis metabolomic profile

The absence of a detectable metabolomic signature in the years preceding MS diagnosis is an important negative finding, despite our modest pre-diagnosis cohort size. While metabolomics has been proposed as a tool for early diagnosis of MS, our data suggest that the broad plasma metabolomic signature of established MS is not yet present a mean of 3.7 years before diagnosis. This suggests that the significant metabolic perturbations associated with MS reflect a consequence of the disease rather than a precursor, and therefore metabolomics is better suited as a biomarker of disease severity and activity rather than a pre-clinical screening tool.

### Strengths and limitations

The principal strengths of this study are its large sample size, rigorous clinical phenotyping, longitudinal follow-up data, inclusion of pre-diagnosis cases, and the use of a standardized metabolomics platform enabling comparability with future studies. The use of a biobank-based control population, while allowing for large-scale comparisons, introduces heterogeneity in health status that was partially addressed by Charlson index adjustment. Controls were slightly older and had a different sex distribution than cases, and residual confounding from comorbidities cannot be excluded. Medication data including usage of lipid-lowering agents was not available. Additionally, although individual BMI data was not available for controls and therefore BMI was not included as a covariate in comparisons between cases and controls, the entire metabolomics cohort is known to have a mean BMI of 28.2 (SD 6.6), which is essentially identical to the MS cohort mean BMI of 28.4 (SD 6.5).

Still, we cannot eliminate the possibility of body composition differences in patients with MS impacting the circulating metabolome. The MRI, NfL, and GFAP subset analyses were limited by small sample sizes and should be considered exploratory.

The effect sizes observed in this study reflect associations with standardized metabolite z-scores rather than absolute concentrations and should be interpreted accordingly. While statistically significant after FDR correction, the modest effect sizes for most associations reflect the inherent heterogeneity of a biobank-based cohort and the multifactorial nature of MS pathophysiology. The clinical relevance of these findings will require validation in independent cohorts. Finally, while the omega-6/omega-3 ratio reflects circulating fatty acid status, it is a proxy for dietary intake rather than a direct measure, and future studies should integrate dietary data to determine whether these associations are primarily driven by dietary patterns or by disease-related metabolic alterations.

## Conclusions

This study provides a comprehensive characterization of the plasma metabolome in MS, identifying coordinated dysregulation of amino acid and lipoprotein metabolism as hallmarks of established disease and disability, and a novel interaction between the omega-6/omega-3 ratio and DMT efficacy as a predictor of future MRI disease activity. These findings support the growing recognition of metabolic dysregulation as a dimension of MS pathophysiology and raise testable hypotheses regarding dietary modification or supplementation as a complement to pharmacological treatment in MS.

## Acknowledgements

This work was funded by the National Multiple Sclerosis Society grant FAN-2407-43752 to R.E. Rodin. B.C. Healy also received funding support from the National Multiple Sclerosis Society. M. Polgar-Turcsanyi and H. Lokhande had no applicable funding sources to report. T. Chitnis received funding support from the National Multiple Sclerosis Society and Department of Defense. We thank the patients who contributed their samples and medical information to research, the Mass General Brigham biobank, and the clinicians who documented clinical visits in the clinical database used for this study.

## Data availability statement

Raw metabolomics data were generated by the Mass General Brigham Biobank in collaboration with Nightingale Health. Anonymized data are available from the corresponding author upon request.

---

**Supplementary Figure 1:** Derivation of metabolite groups by correlation-based hierarchical clustering. (A) Spearman rank correlation matrix of all 162 metabolites, with color indicating the strength and direction of pairwise correlations. (B) Dendrogram from hierarchical clustering of the correlation matrix using a distance metric of (1 – r) and average linkage. The dendrogram was cut at a height corresponding to moderate positive correlations (approximately r = 0.2 – 0.3) to approximate the 30 metabolite groups used in downstream analyses. Manual modifications were made to groups based on known biological pathways.

**Supplementary Figure 2: Metabolite group differences across MS subtypes.** Heatmap of the 30 metabolite groups across MS diagnoses (CIS, PPMS, and SPMS) relative to RRMS as the reference group, derived from linear regression. Cells are colored by regression coefficient. No metabolite group was significantly associated with subtype after false discovery rate correction.

**Supplementary Figure 3: Covariate-adjusted metabolite levels across control, pre-diagnosis, and established MS groups.** Heatmap of covariate-adjusted mean levels for all 30 metabolite groups across 46,443 controls, 25 pre-diagnosis participants, and 411 patients with established MS. Each cell is colored by the covariate-adjusted mean z-score for the corresponding metabolite group and diagnostic group, with the color scale indicating relative metabolite level.

**Supplementary Figure 4: Covariate-adjusted metabolite levels for individual metabolite groups across control, pre-diagnosis, and established MS groups.** Bar charts showing covariate-adjusted mean levels for each of the 30 metabolite groups across 46,443 controls, 25 pre-diagnosis participants, and 411 patients with established MS. Bar height represents the covariate-adjusted mean z-score for each diagnostic group, and error bars represent 95% confidence intervals.

## Supplementary Tables 1-18

Supplementary Table 1. Metabolite list and missingness.

Supplementary Table 2. Metabolite group definitions.

Supplementary Table 3. Individual metabolite associations with MS.

Supplementary Table 4. Metabolite group associations with MS.

Supplementary Table 5. DNEA node-level differential expression.

Supplementary Table 6. DNEA network edges.

Supplementary Table 7. DNEA subnetwork significance (NetGSA).

Supplementary Table 8. DNEA sensitivity analysis.

Supplementary Table 9. Metabolite group associations with EDSS.

Supplementary Table 10. Progressive versus relapsing MS regression.

Supplementary Table 11. Metabolite groups across MS subtypes.

Supplementary Table 12. Time to SPMS conversion.

Supplementary Table 13. Metabolite groups and disease activity.

Supplementary Table 14. Omega-6/omega-3 ratio across six disease activity outcomes.

Supplementary Table 15. Omega-6/omega-3 × DMT efficacy interaction.

Supplementary Table 16. Metabolite groups and MRI biomarkers.

Supplementary Table 17. Pre-diagnosis cohort characteristics.

Supplementary Table 18. Pre-diagnosis metabolite group associations.

