## Supplementary figures and images for "Metabolomics reveals lipid and amino acid signatures of disease severity in multiple sclerosis"

### Figure S1

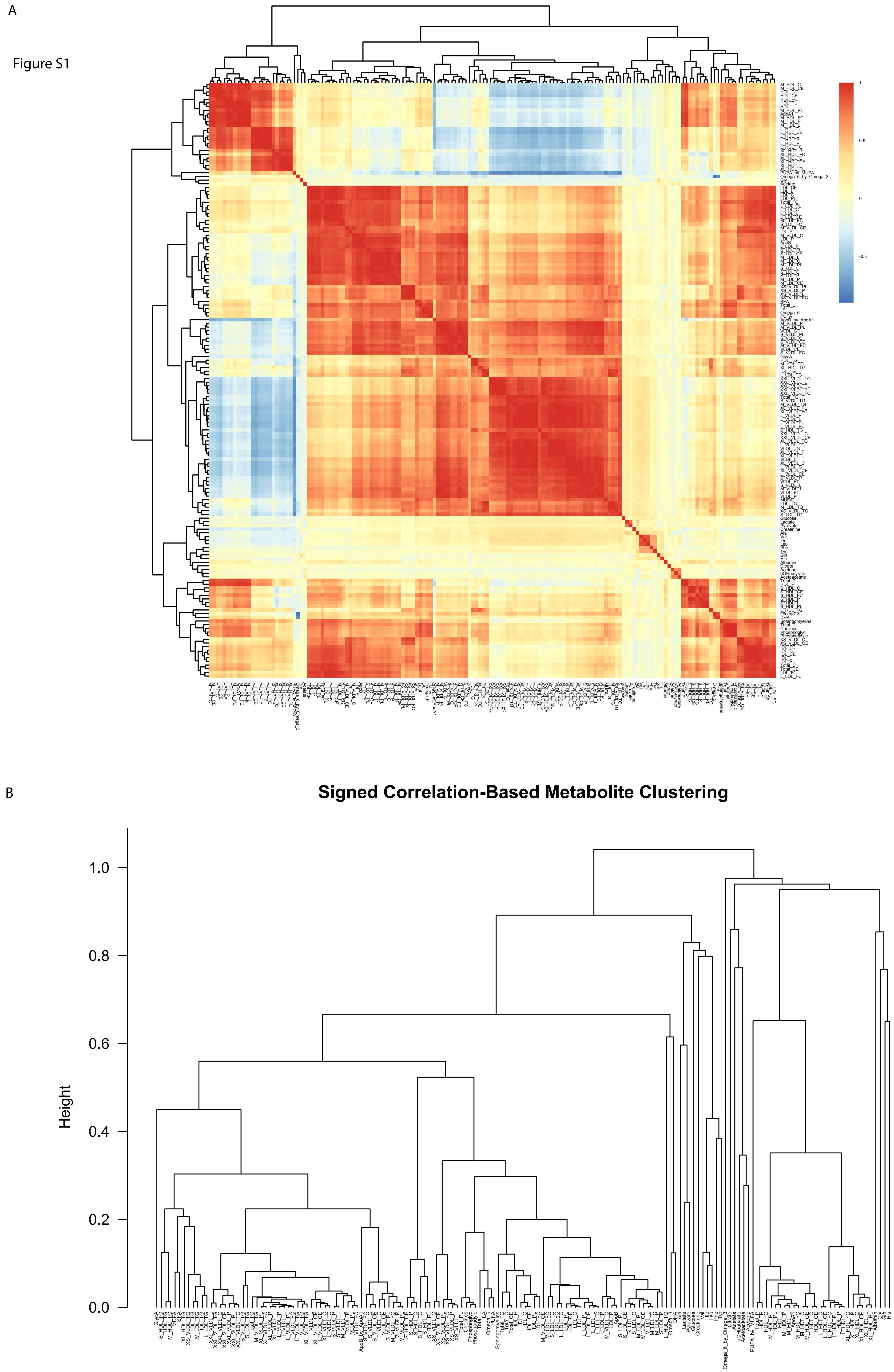

### Figure S2

Figure S2

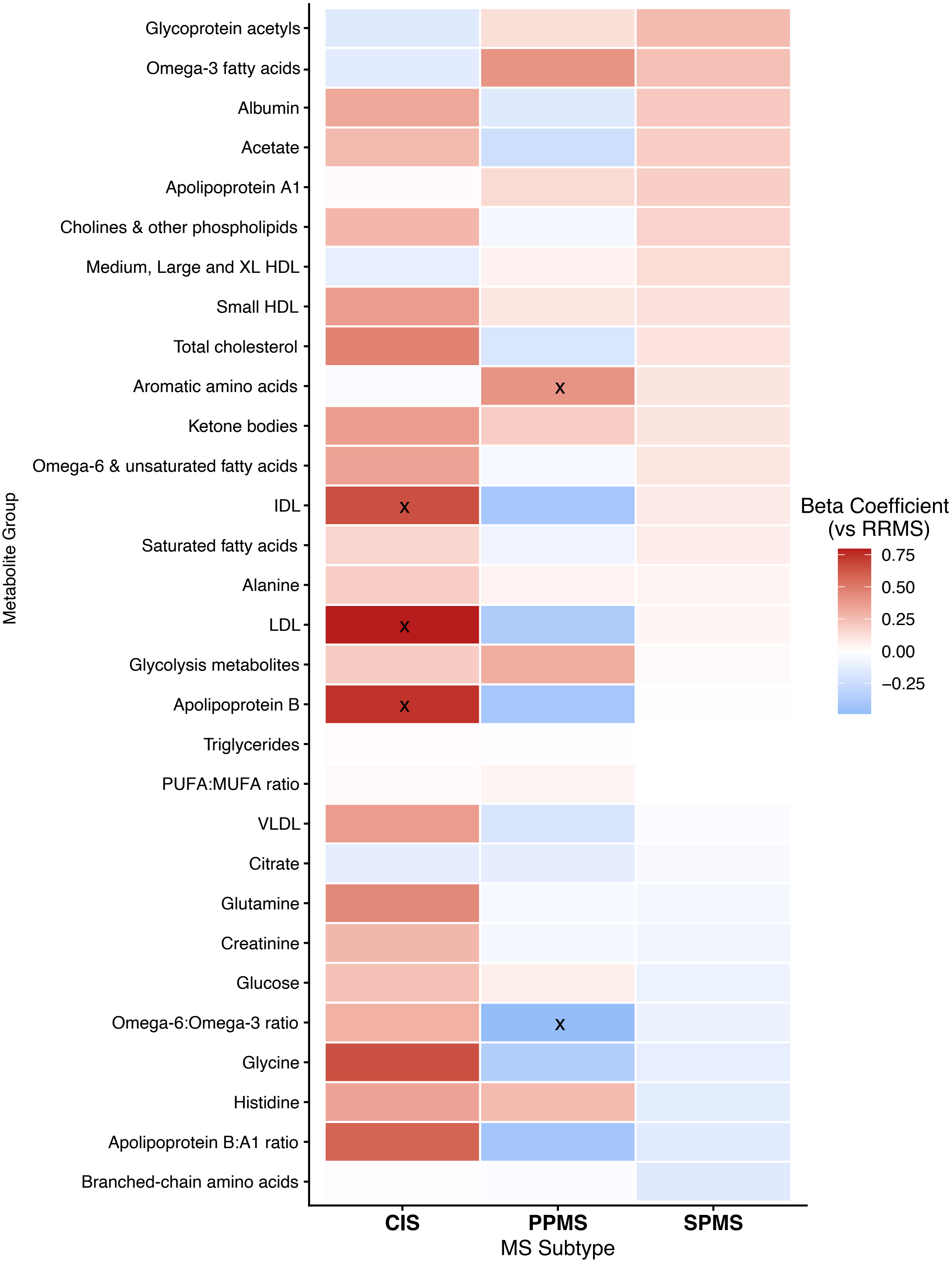


x: $p < 0.05$ (nominal); No FDR significant findings

### Figure S4

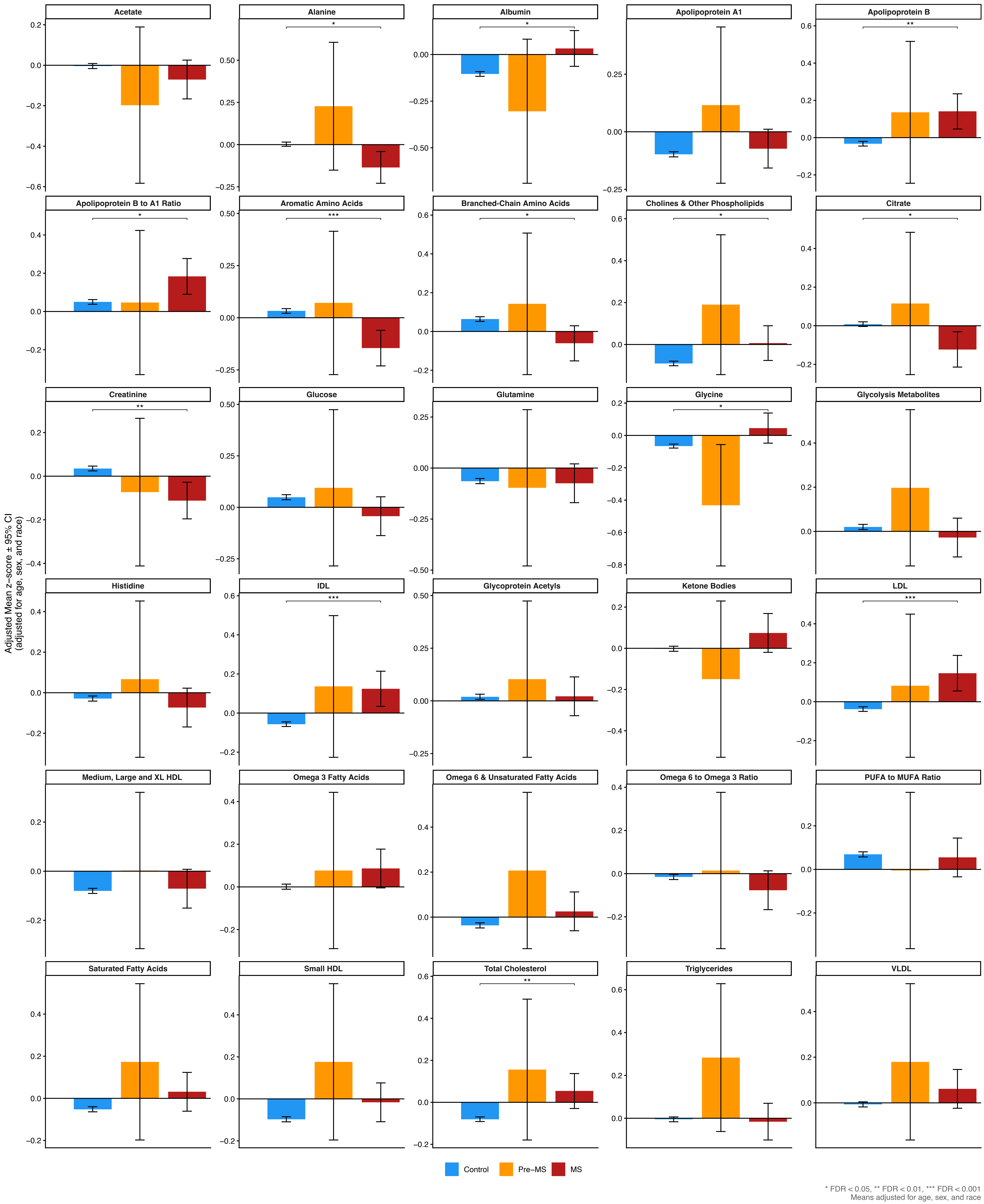
