## Supplementary material for "Metabolomics reveals lipid and amino acid signatures of disease severity in multiple sclerosis": Figure S3

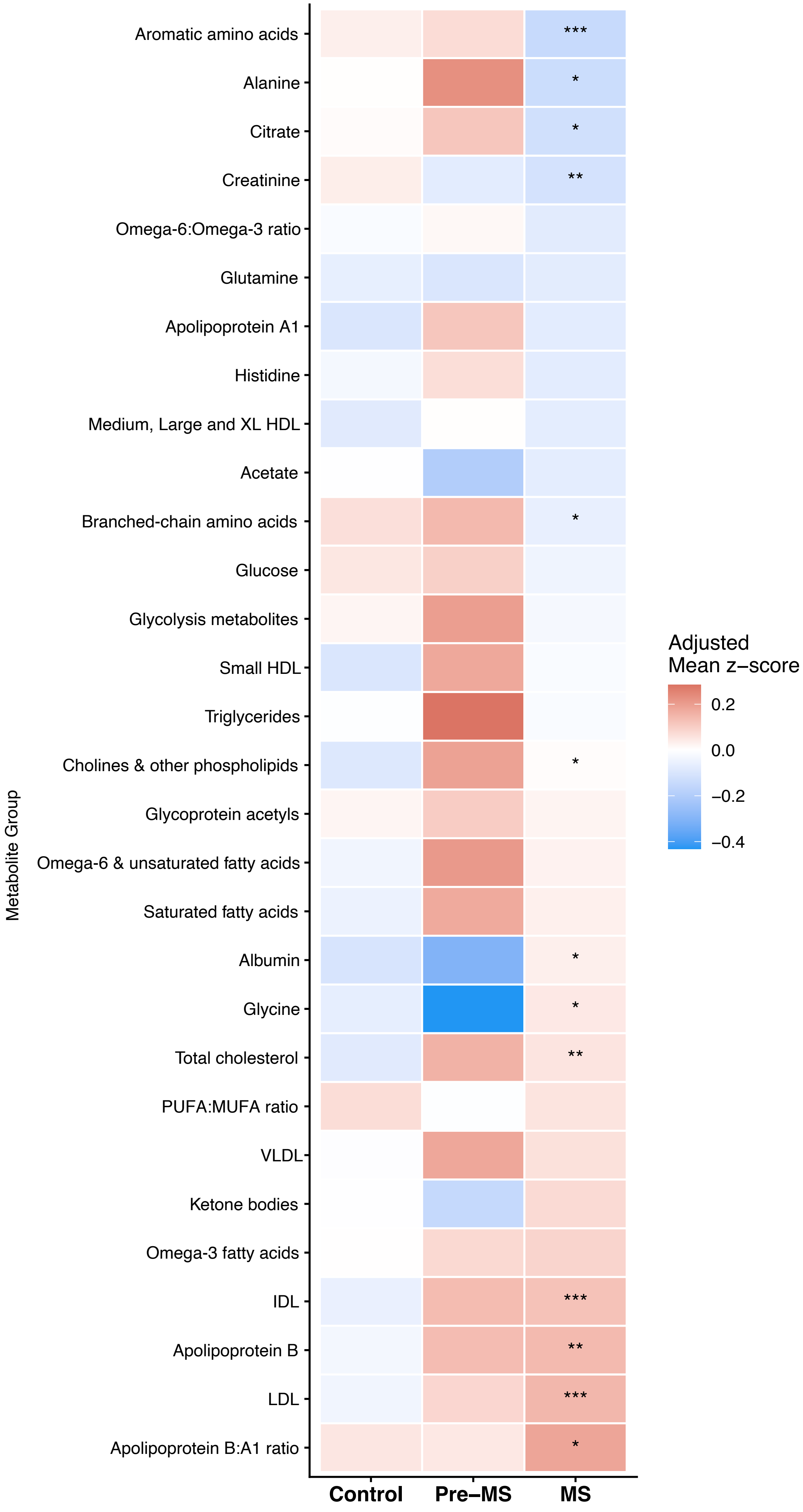


* FDR < 0.05, ** FDR < 0.01, *** FDR < 0.001 vs Non–MS
No significant differences were observed between Pre–MS and Control, or between Pre–MS and MS (all FDR > 0.05)
