## Supplementary Methods for "Metabolomics reveals lipid and amino acid signatures of disease severity in multiple sclerosis"

**Metabolomics quality control**

Quality control consisted of excluding outlier values more than four times the interquartile range from the median of each metabolite, log transforming metabolite values, using a linear regression model to remove the spectrometer effect, and scaling residuals to a mean of zero and standard deviation of one for each metabolite. Of the 249 metabolite values delivered by the Nightingale platform, 168 are absolute levels and 81 are derived ratios. To minimize redundancy, metabolites representing the sum of other included metabolites were removed (for example, individual branched-chain amino acids were retained and total branched-chain amino acids removed), as were nearly all derived ratios except for key ratios of biologic interest. After filtering, 162 unique metabolites remained for downstream analyses.

**Clinical phenotyping details**

Relapses and MRI changes were identified through Oracle database review and/or medical record review. A relapse was defined as new or worsening neurological symptoms attributable to MS, lasting at least 24 hours, in the absence of fever, infection, or other acute medical illness, occurring at least 30 days after the onset of any prior relapse, and documented by a treating neurologist. MRI activity was defined as the presence of new or enlarging T2 hyperintense lesions or gadolinium-enhancing lesions on brain or spinal cord MRI, as documented in the radiology report. Twelve patients were excluded from relapse analysis due to incomplete relapse data. There were 95 relapse events and 99 MRI activity events amongst 112 cases, and not all patients with clinical relapses underwent concurrent MRI. The high amount of disease activity in this cohort reflects >50% of the patients being on no disease-modifying therapy (DMT) or low-efficacy DMT when samples were collected.

DMT at the time of sample collection was extracted from the Oracle database or chart review. DMTs were classified as low-efficacy (glatiramer acetate, interferon, long-term corticosteroids, mycophenolate), moderate-efficacy (sphingosine-1-phosphate receptor modulators, fumarates, cyclophosphamide, teriflunomide), and high-efficacy (B-cell depletion therapy, alemtuzumab, natalizumab). For participants with MS, body mass index (BMI) was extracted from the clinical visit closest to the biobank sample collection date.

**Imputation of missing values**

Across the full dataset there was very low metabolite data missingness (0.31%; Supplementary Table 1). Missing value imputation was performed separately within MS and control groups using K-nearest neighbor (KNN) imputation (*k* = 10) from the Bioconductor impute package. Group-specific imputation was used to prevent cross-group biological contamination of imputed values.

**Derivation of metabolite groups**

To reduce dimensionality and comprehensively assess biological pathways, metabolites were clustered into groups for downstream analyses. Pairwise associations between all 162 metabolites were assessed using Spearman rank correlation coefficients. A metabolite–metabolite correlation matrix was constructed (Supplementary Fig. 1A). To identify groups of co-varying metabolites, hierarchical clustering was performed on the correlation matrix using a distance metric defined as (1 - r), where (r) is the Spearman correlation coefficient. Clustering was conducted using average linkage. The resulting dendrogram was cut at a height corresponding to moderate positive correlations (approximately *r* = 0.2-0.3), based on inspection of the dendrogram and cluster stability, to define discrete metabolite groups (Supplementary Fig. 1B).

To enhance interpretability, data-driven clusters were reviewed in the context of established metabolic pathways. Where appropriate, minor refinements were made to ensure biological coherence of groups while maintaining the overall correlation structure. In total, 30 groups were identified ranging in size from 1 to 42 metabolites (Supplementary Table 3). All downstream analyses of metabolite groups used the mean z-score across metabolites within each of the 30 groups.

**Differential network enrichment analysis**

Prior to DNEA, metabolite values were residualized against age, sex, race, and Charlson comorbidity index across the full cohort. The regularization parameter lambda was optimized using the Bayesian Information Criterion, yielding an optimal lambda of 0.003192. Stability selection was performed with 500 replicates and sub-sampling to account for the imbalance in sample sizes and to identify robust edges. Network clustering was performed using the consensus clustering approach implemented in DNEA.
